# Impacts of radiation on the bacterial and fungal microbiome of small mammals in the Chernobyl Exclusion Zone

**DOI:** 10.1101/2020.05.28.120717

**Authors:** Rachael E. Antwis, Nicholas A. Beresford, Joseph A. Jackson, Ross Fawkes, Catherine L. Barnett, Elaine Potter, Lee Walker, Sergey Gaschak, Michael D. Wood

## Abstract

Environmental impacts of the 1986 Chernobyl Nuclear Power Plant accident are much debated, but the effects of radiation on host microbiomes has received little attention to date. We present the first analysis of small mammal gut microbiome from the Chernobyl Exclusion Zone in relation to total absorbed dose rate and including caecum as well as faeces sample. The associations between microbiome communities and radiation exposure varied between host species. Associations between microbiome and radiation was different for analyses based on ambient versus total weighted absorbed dose rates. We found considerable variation between patterns for faecal and gut samples of bank voles, suggesting faecal samples are not an accurate indicator of gut composition. For bank vole guts, associations between radiation and bacterial community composition were robust against geographical and habitat variation. We found limited associations between radiation and fungal communities. Host physiological mechanisms or environmental factors may be driving these patterns.

## INTRODUCTION

Multicellular organisms host a complex community of microbes (the microbiome) that is critical for host health and function ^1,2^. The gut microbiota has been shown to affect animal development, immune response, food digestion and behaviour ^3^. Microbiome composition varies according to biological and environmental factors such as host species ^4^, host age ^5^, diet ^6^, season ^7^, and contaminant-induced stress ^8^, among others ^9^. Less well understood is the relationship between radiation exposure and microbiome composition, particularly in wild animal systems.

Over the last decade there has been a growing interest in the effect of contaminants on the composition of the gut microbiome, with some studies reporting changes in the two most prevalent bacterial phyla within the gut ^10,11^, namely Firmicutes and Bacteroidetes ^12^. Different chemical stressors have been found to affect Firmicutes: Bacterioidetes (F:B) ratios, with As ^13^, Cd ^14^, chlorpyrifos ^15^, permethrin ^16^ and pentachlorophenol ^17^ leading to decreases in F:B ratios, whereas Pb ^11^ and carbendazim ^18^ cause increased F:B ratios. These changes in gut microbiota composition have also been linked to changes in host immune responses ^12^.

High acute radiation exposure (> 1 Gy) has been shown to influence gut microbial communities (e.g. Dubois & Walker 1989; Packey & Ciorba 2011), leading to the suggestion that gut microbiota could be a potential biomarker of radiation exposure ^21,22^. Improvements in the responses of both humans and model organisms to acute radiation exposure have also been observed when bacterial probiotics (particularly *Lactobacillus* spp.) were administered (e.g. Demers *et al.* 2014; Meng-Meng *et al.* 2017). Some studies suggest that faecal microbiomes may be associated with lower radiation exposure in contaminated environments, such as the Chernobyl Exclusion Zone (CEZ) ^25,26^. For example, Lavrinienko et al. ^25^ report a reduction in the faecal F:B ratios of bank voles (*Myodes glareolus*) at their most contaminated sites (mean ambient dose rate 30 μSv h^−1^). Radiation-induced changes in the microbiome of skin and feathers in organisms from Chernobyl have also been investigated. Lavrinienko et al. ^26^ found no effect of radiation on skin microbiome of bank voles, but radiation-induced changes in feather baterial communities have been suggested ^27,28^.

The extent to which radiation exposure is affecting wildlife in Chernobyl is highly contested ^29,30^. A fundamental problem with many of the studies undertaken to date is that they use ambient dose rates from the air (often reported in units of absorbed radiation dose rate for humans, μSv/h), rather than estimating the total abosrbed dose rate of study organisms, taking account of both internal and external exposure ^31^. As such, it is not possible to accurately determine dose-effect relationships, making interpretation of these studies difficult. Here we present the first study of GI tract microbiome composition in CEZ small mammals for which individual total absorbed dose rates have been estimated. Previous studies in the CEZ have only considered the bacterial microbiome of one small mammal species (bank vole) using faecal samples; here we report on the faecal microbiome of four small mammal species using faecal samples, as well as the first direct analysis of gut microbiome using caecum samples from bank voles. In addition, our microbiome analysis includes both bacteria and fungi, extending our limited general knowledge on the fungal component of animal microbiomes.

## METHODS

### Field sampling in the Red Forest (2017)

In August 2017, we sampled small mammals from the Red Forest, an area of c. 4-6 km^2^ over which pine trees were killed by radiation in 1986; subsequently there has been sparse regrowth of deciduous trees and some understorey vegetation. In 2016, approximately 80% of the Red Forest was damaged by fire. Our 2017 sampling sites (Figure 1) included a total of eight sites across three burn categories, namely ‘burnt with regrowth’ (n = 2), ‘burnt with minimal regrowth’ (n = 3) and ‘unburnt’ (n = 3). At each of these sampling sites, a 60 m × 60 m trapping grid was used, with traps positioned at 10 m intervals (each grid comprised a total of 49 traps). To maximise trapping success, the trapping grids were established one week prior to the beginning of the study and pre-baited with rolled oats and carrots/cucumber. Trapping occurred over eight consecutive days; traps were baited and set each evening and visited early in the morning to retrieve captured small mammals. The small mammals were transferred to the Chernobyl field station where each animal was live monitored to determine its ^137^Cs whole body activity concentration using an unshielded 51 mm × 51 mm NaI (Tl) detector (GMS 310 core gamma logger) supplied by John Caunt Scientific Ltd. Additional regular background measurements were made each day. The detector was calibrated using the results for small mammals (n = 14) that were live monitored on the GMS 310 and subsequently analysed using a calibrated detector at the Chornobyl Center’s main laboratory (R^2^ = 0.98). The limit of detection (LOD) was estimated as three times the standard deviation of the background measurement. The sex of each animal was determined and their live mass recorded.

**Figure 1.**
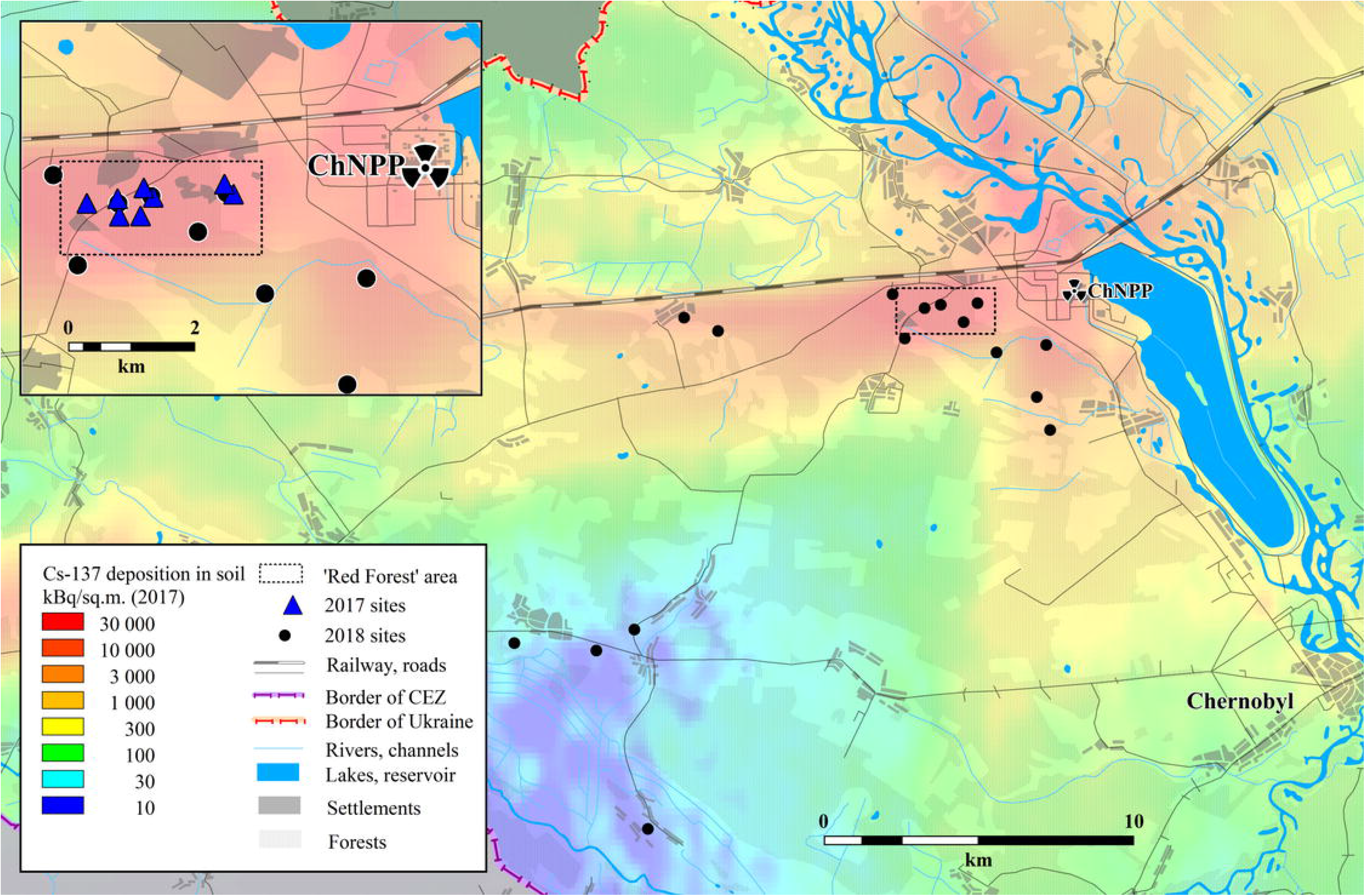
Location of the study sites in the CEZ where small mammals were trapped in 2017 and 2018; the approximate location of the Red Forest is indicated by the black rectangle. The underlying 137Cs soil data shown (decay corrected to summer 2017) are from Shestopalov (1996).

Freshly excreted faecal samples were collected directly from animals for subsequent microbiome analysis. We sampled striped field mice (*Apodemus agrarius*; n = 29), yellow-necked mice (*Apodemus flavicollis*; n = 58), wood mice (*Apodemus sylvaticus*; n = 27) and bank voles (*Myodes glareolus*; n = 22; Table S1). Faecal samples were immediately placed into vials containing 100% ethanol and subsequently stored at −20°C. Samples were transported under licence to the University of Salford (UK); sample integrity was maintained during transit using dry ice and the samples were then stored at −20°C prior to DNA extraction. We used fur clipping to mark each small mammal prior to release at the point of capture, which allowed us to check whether each capture over the subsequent days was a new animal. Only faeces from new animal captures were included in this study.

### Field sampling across the CEZ (2018)

Small mammals were trapped in July/August 2018 over 10 consecutive days, with only bank voles included in this study. Twelve transects of Sherman traps were established at sites across a gradient of ambient dose rates (Figure 1). Each transect measured 290 m with a trap interval of 10 m (30 traps per transect). The 2018 sampling adopted the same protocol for baiting and collection of captured animals that was used in 2017. For some of the analyses, bank voles from 2018 have been categorised by collection ‘site category’, defined as inside or outside of the Red Forest.

Captured animals were transferred to the Chernobyl field station, where each animal was live monitored to quantify the whole-body activity concentrations of both ^137^Cs and ^90^Sr. This was done using a new field portable Radioanalysis of Small Samples (ROSS) detector developed at the University of Salford ^32^. ROSS comprises a sample holding chamber with a capacity of 170 × 60 × 50 mm. Two CsI gamma detectors (each measuring 70 × 40 × 25 mm) were mounted on opposite sides of the sample holding chamber and two plastic scintillator beta detectors were mounted one above (100 × 50 × 0.5 mm) and one below (100 × 60 × 0.5 mm) the chamber. The entire assembly was enclosed within a lead shield (>10 mm thickness). ROSS was calibrated using ^137^Cs and ^90^Sr standards developed by Chornobyl Center; standards ranged from 4 to 20 g to represent small mammals. We included Cs-only standards, Sr-only standards and mixed standards. Counting of the Cs standards on the beta detectors provided a correction for the influence of gamma emissions. Multiple background counts were performed daily (at least nine per day) and the LOD was estimated using the method described by Currie ^33^.

Bank voles were killed by an overdose of anaesthetic (isofluorane) followed by exsanguination (in accordance with Schedule 1 of the Animals (Scientific Procedures) Act 1986). The sex and mass of each animal was recorded. Gastrointestinal tracts (n = 142) were dissected immediately and stored in laboratory vials containing 100% ethanol and stored at −20°C. The frozen vials were transported to the University of Salford under licence and stored as described for faeces.

### Dosimetry: Ambient dose rate

All dose data are provided as supplementary data. At every trapping location in 2017 and 2018, ambient dose rate (μSv h^−1^) was measured using an MKS-01R meter at 5 cm above the soil surface.

### Dosimetry: Estimation of small mammal total absorbed dose rate for the 2017 study

Soil samples (0 - 10 cm soil depth) were available from each of the trapping sites used in the 2017 study. These samples were analysed using laboratory detectors at Chornobyl Centre to determine ^137^Cs and ^90^Sr activity concentrations within the soil (see ^34^ for methodology). For each small mammal species, an external dose conversion coefficient was calculated using the ERICA Tool version 1.2 ^35^. To define the geometry for each species, the length, width and height were determined through literature review. Soil activity concentrations were input into the ERICA Tool and external dose rates estimated using the derived external dose conversion coefficients and appropriate occupancy factors (assuming 50% of time in soil and 50% on the soil surface for mice and 70% in soil and 30% on soil for bank voles).

The measured ^137^Cs whole body activity concentrations were used to determine the internal absorbed dose from ^137^Cs. In 2017. The internal ^90^Sr activity concentrations were not directly measured; these were estimated using the species-specific transfer parameters measured in the CEZ and presented by Beresford et al. ^34^ and the soil ^90^Sr activity concentrations for the appropriate sampling site. For each small mammal species, an internal dose conversion coefficient was calculated using the ERICA Tool and the same assumed geometries as used for the external dose conversion coefficient derivation. The ERICA Tool was then run using the default radiation weighting factors to calculate total weighted absorbed dose rate. Whilst other radionuclides (e.g. Pu-isotopes and ^241^Am) are present in the CEZ, Beresford et al. ^34^ demonstrated that the contribution of these isotopes to small mammal total absorbed dose rate of small mammals within the Red Forest was low (< 10 %).

For each individual animal, the total weighted absorbed dose rate (hereafter referred to as the total absorbed dose rate) was calculated by summing the internal and external dose rates for that individual.

### Dosimetry: Estimation of small mammal total absorbed dose rate for the 2018 study

Soil activity concentrations were not available for all of the sites within the 2018 study. However, Beresford et al. ^34,36^ reported relatively good agreement between estimated external dose from ^137^Cs and the external ambient dose field at sites in the CEZ. Based on our small mammal dose rate data from 2017, the mean ratio of external dose from ^137^Cs to the external ambient dose field is 0.98. We used the dosimetry approach of the ERICA Tool, which assumes a shielding effect from fur and skin for external beta exposure ^37^ and subsequently also adopted by the ICRP ^38^. The estimated contribution of ^90^Sr (a beta emitter) to the external whole-body dose rate of small mammals is therefore negligible. Therefore, the external dose rates measured at each trapping location were used to estimate the external absorbed dose rate for each small mammal using the occupancy factors defined above.

The ^137^Cs and ^90^Sr whole body activity concentrations measured using ROSS were input into the ERICA Tool and the species-specific internal dose conversion coefficients were used to estimate the internal absorbed dose rate for each animal. At the lowest contamination sites in 2018, some of the whole-body activity concentrations for both ^137^Cs and ^90^Sr were below the LOD. Using these LOD values to determine total absorbed internal dose rate led to a maximum estimated dose of 0.6μGy h^−1^, introducing some uncertainty in radiation exposure estimates at the lowest end of our total absorbed dose rate range.

### Dosimetry: Incorporation of estimated dose rates with subsequent analyses

We assigned animals to total absorbed dose rate categories based on the suggested derived consideration reference level for ICRPs Reference Rat ^38^, i.e. approximately 4-42 μGy h^−1^. As such, animals estimated to receive total absorbed dose rates of <4 μGy h^−1^ were assigned ‘low’, those with estimated dose rates of 4-42 μGy h^−1^ assigned ‘medium’, and those >42 μGy h^−1^ assigned to a ‘high’ category. The ‘high’ and ‘low’ total absorbed dose rates are in-effect also a comparison of inside and outside the Red Forest (i.e. the ‘inside’ and ‘outside’ Red Forest site categories; Table S1).

We correlated ambient and total estimated absorbed dose rates using a Spearman’s rank correlation, To quantify whether correlation coefficients varied based on radiation dose, we also repeated the correlations for each total absorbed dose rate category separately, and visualised these using a scatter plot in ggplot2 ^39^.

### DNA extraction and molecular work

For faecal samples, we extracted DNA from the full sample (~0.1 g) of the four host species. For gut samples, we isolated ~25% of the distal end of the caecum of bank voles and homogenised the contents by hand in sterile petri dishes, before weighing out ~0.1 g for DNA extraction. We conducted all DNA extractions using the PureLink™ Microbiome DNA Purification Kit (Invitrogen, UK) according to the manufacturer’s instructions.

To identify bacterial communities, we conducted 16S rRNA amplicon sequencing according to Kozich et al. ^40^ and Griffiths et al. ^41^. Briefly, we ran PCRs in duplicate using Solis BioDyne 5x HOT FIREPol^®^ Blend Master Mix, 2μM primers and 15ng of sample DNA under thermocycling conditions of 95 °C for 10 min; 25 cycles of 95°C for 30s, 55°C for 20s, and 72°C for 30s; and a final extension of 72 °C for 8 minutes. We included negative (extraction blanks) and positive (mock community) controls. We combined PCR replicates into a single PCR plate and cleaned these using HighPrep™ PCR clean up beads (MagBio, USA) according to the manufacturers’ instructions. We quality checked PCR products throughout on an Agilent 2200 TapeStation. To quantify the number of sequencing reads per sample, we constructed a library pool using 1ul of each sample. We conducted a titration sequencing run using this pool with a v2 nano cartridge (2 × 150bp) on the Illumina MiSeq platform. We calculated the volume of each sample required based on the percentage of reads obtained and pooled these accordingly. We sequenced the final normalised library using paired end (2 × 250bp) reads on a v2 cartridge on an Illumina MiSeq at the University of Salford.

We identified fungal communities via the ITS1F-2 rRNA gene using a modified protocol of Smith & Peay ^42^ and Nguyen et al. ^43^, as in Griffiths et al. ^44^. We ran PCRs in duplicate using thermocycling conditions of 95°C for 10 minutes, followed by 28 cycles of 95°C for 30s, 54°C for 45s, and 72°C for 60s; with a final extension at 72°C for 10 minutes. We included extraction blanks and a mock community as negative and positive controls, respectively. We quantified and normalised individual libraries as described above, before conducting full paired-end sequencing using Illumina v2 (2 × 250bp) chemistry on an Illumina MiSeq at the University of Salford.

### Pre-processing of amplicon sequencing data

Unless otherwise stated, we conducted all data processing and analysis in RStudio (v1.2.1335) ^45^ for R (v3.6.0) ^46^. A total of 13,371,018 raw sequence reads from 279 samples, plus one mock community and ten negative controls, were generated during 16S rRNA gene amplicon sequencing, which we processed in DADA2 v1.5.0 ^47^. Modal contig length was 253bp once paired-end reads were merged. We removed sequence variants (SVs) with length >260bp (26 SVs; 0.002% of total sequences) along with chimeras and five SVs found in the negative controls. We assigned taxonomy using the SILVA v132 database ^48,49^. DADA2 identified 20 unique sequence variants in the sequenced mock community sample comprising 20 bacterial isolates. We stripped out mitochondria from samples along with SVs with less than 0.0001% abundance across all samples. We removed three samples from which poor sequence data were obtained (<1000 reads), leaving a median of 26,106 reads (6,557 to 119,871) per sample. We exported the final SV table, taxonomy table and sample metadata to the phyloseq package ^50^.

We obtained a total of 2,778,887 raw sequence reads from 279 samples (plus mock community and negative controls) during ITS rRNA gene sequencing. We trimmed remaining adapters and primers from ITS rRNA sequencing data using cutadapt ^51^ in RStudio. As with 16S rRNA sequence data, we pre-processed ITS rRNA amplicons in DADA2 v1.5.0 ^47^. Modal contig length was 219bp (167-457bp) once paired-end reads were merged. We did not conduct additional trimming based on sequence length as the ITS region is highly variable ^52^. We removed chimeras and one contaminant found in the negative controls, and then assigned taxonomy using the UNITE v7.2 database ^53^. DADA2 identified 12 unique sequence variants in the sequenced mock community sample comprising 12 fungal isolates. We removed 54 samples from which poor sequence data were obtained (<500 reads), leaving a median of 1863 reads (506 to 17,226) per sample. As with 16S rRNA data, we exported the final SV table, taxonomy table and sample metadata to the phyloseq package ^50^ for subsequent analysis.

### Community analyses

For both bacterial and fungal community data, we normalised the clean count data using centred-log ratio (clr) transformations ^54^ in phyloseq ^50^, and visualised beta-diversity (between species and between sample type, i.e. gut or faeces) using PCA plots with Euclidean distances in ggplot2 ^39^. We used PERMANOVAs to test for significant differences in beta-diversity according to species, sex, sample type and total absorbed dose rate category using the adonis function in the vegan package ^55^. We agglomerated the data to family level and visualised differences in clr-transformed data according to the five sampling groups (faecal samples from the three mice species, plus faecal and gut samples from bank voles) using jitter box plots in ggplot2 ^39^. We tested for significant differences between sampling groups in the clr-transformed values of these 24 families (12 per microbial kingdom) using Kruskal-Wallis non-parametric tests with Dunn’s pairwise tests and Hochberg-adjusted p values in the dunn.test and FSA packages ^56,57^. We also converted the raw SV counts to relative abundance and visualised the 12 most abundant families (for each kingdom separately) as a stacked chart according to species and sample type.

We then split the clr-transformed data by sampling year and re-ran the PERMANOVA analysis for the 2017 faecal samples with species, sex, total absorbed dose rate category, grid line and burn category as predictor variables. We visualised the variation in clr-transformed values for the 12 most abundant genera in faecal samples from yellow-necked mice (as this was the only species with sufficient samples across all three burn categories; Table S1) using PCA plots of beta-diversity and jitter plots for the clr values of the 12 most abundant genera. We also re-ran the PERMANOVA analysis for the 2018 gut data with total absorbed dose rate category, site category, sex and transect line as predictor variables, and again visualised these using PCA plots of beta-diversity and jitter plots of the clr values of the 12 most abundant genera for each microbial kingdom separately.

To determine whether microbiome beta-diversity correlated with total absorbed dose rate independently to geographic location for bank vole gut data, we conducted partial Mantel tests using the vegan package ^55^ on the gut data from bank voles. We constructed microbiome distance matrices in phyloseq ^50^ between samples for bacteria and fungi separately using Euclidean distances of clr-transformed data. We generated Euclidean distance matrices from the total absorbed dose rate data for each individual using the proxy package ^58^. Finally, we constructed a geographic distance matrix between radiation distance and samples using longitude and latitude coordinates in Microsoft Excel. We then ran partial Mantel tests with Spearman’s rank correlation between total absorbed dose rate distance and microbiome distance (for bacteria and fungi separately), with the geographic distance matrix as a covariate.

For bank vole gut data, we calculated alpha diversity (SV richness, and community evenness using the inverse Simpson index) of bacterial and fungal communities by subsampling the raw SV count table to a standardised number of reads (equal to the sample with the lowest number of reads) using an iterative approach (100 times), and averaging the diversity estimates across these iterations. We correlated these alpha-diversity measures with total absorbed dose rate for each bank vole sample using Spearman’s correlations.

We used Spearman’s rank correlation (with Benjamini-Hochberg corrected p values and False Discovery Rate adjustment) in the associate function of the microbiome package ^59^ to identify relationships between the two radiation dose measures (ambient and total) and clr-transformed 16S and ITS rRNA sequence data, agglomerated to genus level. These analyses were conducted separately for the gut and faecal samples, according to host species. We then visualised the resultant correlation coefficients using heatmaps in ggplot2 ^39^.

We calculated Firmicutes: Bacteriodetes (F:B) ratios in vole guts using both clr-transformed data, and data converted to relative abundance (as in ^25^). We also calculated F:B ratios in faecal samples of all four mammal species using relative abundance data. We visualised these ratios according to total absorbed dose rate category using jitter plots. We tested for significant differences between total absorbed dose rate categories within a set of data using Kruskal-Wallis non-parametric tests, with Dunn’s pairwise tests and Hochberg-adjusted p values where necessary.

## RESULTS

### How do ambient dose rates compare to total absorbed dose rates?

There was a significant positive correlation between ambient and total absorbed dose rates across all animals captured during both 2017 and 2018 (r = 0.529, p < 0.001), although there was considerable variation around the data (Figure 2). When data were split into the three different total absorbed dose rate categories, all relationships remained statistically significant (low (< 4 μGy h^−1^): r = 0.898, p < 0.001; medium (4-42 μGy h^−1^): r = 0.879, p < 0.001; high (>42 μGy h^−1^): r = 0.236, p < 0.001), however the variation in the ‘high’ dose data was particularly evident (although note the greater sample size; Figure 2). The estimated total absorbed dose rate gives a better estimation than ambient dose rate of each individual’s radiation exposure, and hence we used total absorbed dose rates for the majority of our analyses. In this study we have used the ERICA dosimetry approach, which assumes shielding by fur and skin of beta radiation and consequently the external dose rates from ^90^Sr are estimated to be negligible. We acknowledge that estimates using different modelling approaches may lead to a higher estimated external dose rate from ^90^Sr (e.g. ^60^). We have estimated the external dose contributions using an alternative model (https://wiki.ceh.ac.uk/x/9wHbBg) which does not consider fur or skin shielding ^61,62^. Using this model, we find that the maximum difference in total dose rate estimate would be approximately 30%, with an average difference of about 10%.

**Figure 2.**
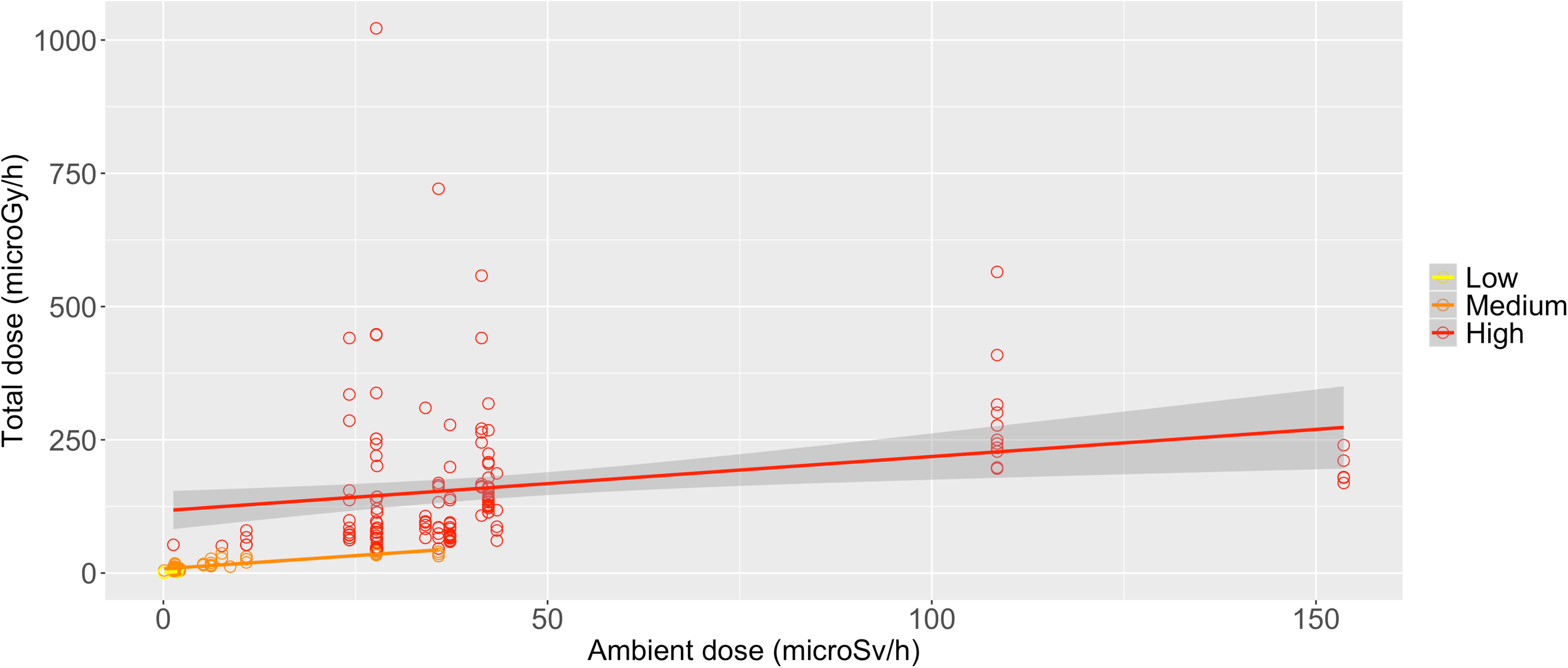
Relationship between ambient and total absorbed dose rates associated with animals sampled in the study, split into total absorbed dose rate categories (low <4 microGy/h; medium = 4-42 microGy/h; high > 42 microGy/h).

### How does microbiome beta-diversity vary according to host factors and total absorbed dose rate?

A PERMANOVA analysis demonstrated sample type (F_1,260_ = 49.408, R^2^ = 0.143, p = 0.001), species (F_3,260_ = 9.846, R^2^ = 0.085, p = 0.001) and total absorbed dose rate category (F_2,260_ = 2.631, R^2^ = 0.015, p = 0.001) all significantly predicted bacterial community beta-diversity, whereas sex did not (F_1,268_ = 1.206, R^2^ = 0.003, p = 0.138). Similarly, fungal community beta-diversity was significantly predicted by sample type (F_1,212_ = 12.574, R^2^ = 0.052, p = 0.001), species (F_3,212_ = 2.183, R^2^ = 0.027, p = 0.001), and total absorbed dose rate category (F_2,212_ = 2.024, R^2^ = 0.016, p = 0.001), but not sex (F_2,212_ = 1.108, R^2^ = 0.004, p = 0.206). As such, sample type was the biggest driver of variation in both bacterial and fungal community composition. Differences between host species were much more evident for bacterial community composition than for fungal community composition (Figures 3a, b), for which 8.5% and 2.7% of the variation was explained by host species, respectively. A full description (with statistical testing) of the differences in community composition based on host species and sample type can be found in Supplementary Material.

**Figure 3.**
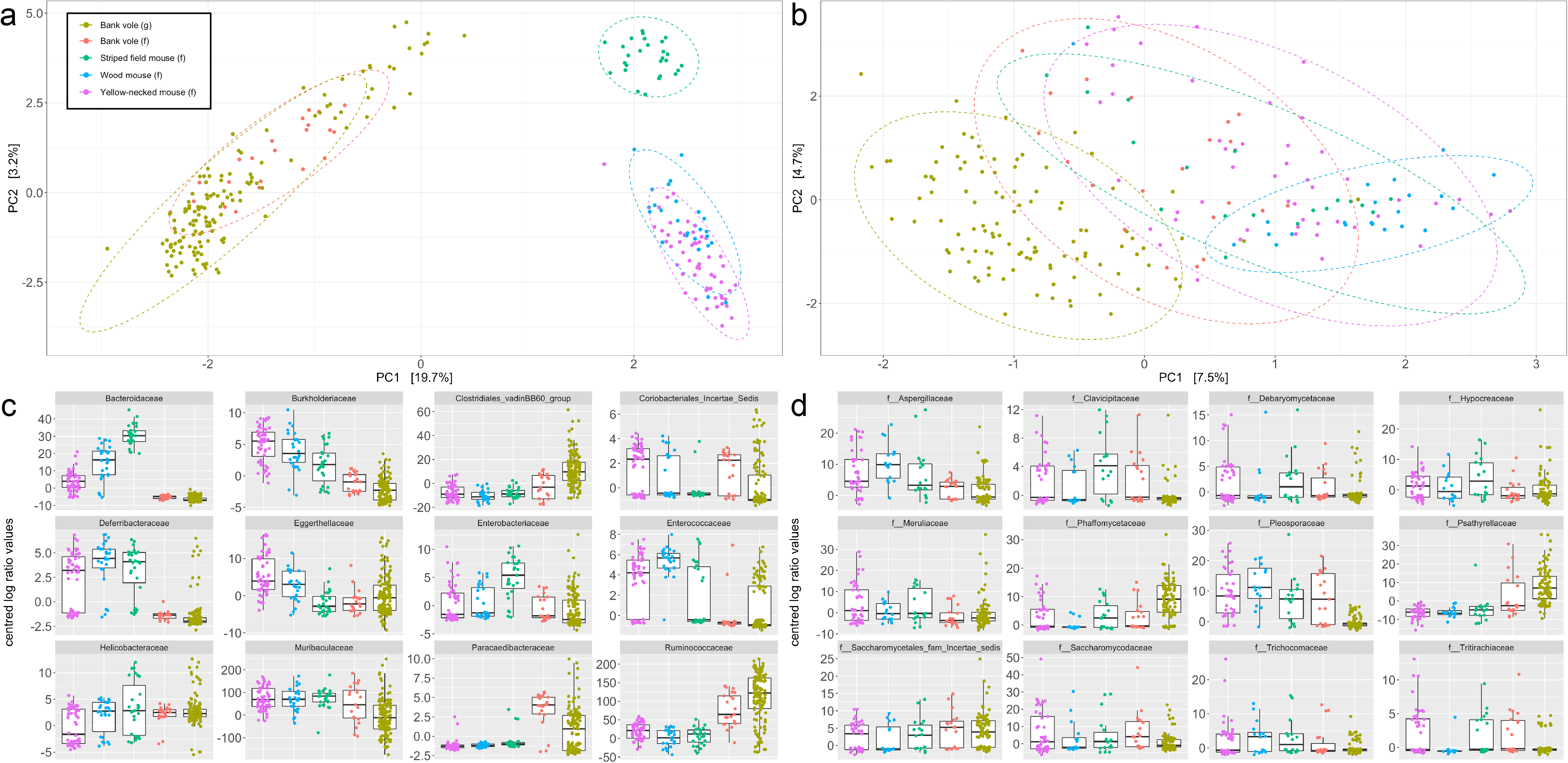
PCA plots showing Euclidean distances of clr-transformed bacterial (a) and fungal (b) communities associated with faecal and gut samples from four small mammal species in the Chernobyl Exclusion Zone. Jitter plots displaying the clr values of the 12 most abundant bacterial (c) and fungal (d) families across the five sampling groups (faecal samples for the three mice species along with faecal and gut samples for the bank voles).

### How does beta-diversity of faecal samples vary according to host species, burn category and total absorbed dose rate?

When using faecal samples (i.e. 2017 data) only, the PERMANOVA analysis indicated that host species (F_3,128_ = 11.944, R^2^ = 0.217, p = 0.001; Figure S3), burn category (F_2,128_ = 1.632, R^2^ = 0.020, p = 0.004; Figure S3) and grid line (F_5,128_ = 1.562, R^2^ = 0.047, p = 0.001) had a significant effect on beta-diversity of faecal bacterial communities, but that total absorbed dose rate category (F_1,128_ = 0.912, R^2^ = 0.006, p = 0.616) and sex (F_1,128_ = 1.103, R^2^ = 0.007, p = 0.228) did not. There were only sufficient samples for yellow-necked mice to visualise differences in microbiome composition across all three burn categories (Table S1). Although the differences were relatively small in the 12 most abundant bacterial genera, some showed directional changes based on burn category (Figure S4), for instance, Bacteriodes were most abundant at ‘burnt (minimal regrowth)’ sites and least abundant at ‘unburnt’ sites, whereas Ruminococcaceae_UCG-003 showed the inverse.

As with bacterial communities, host species (F_3,111_ = 1.945, R^2^ = 0.048, p = 0.001), burn category (F_2,111_ = 1.969, R^2^ = 0.033, p = 0.001) and grid line (F_5,111_ = 1.726, R^2^ = 0.072, p = 0.001) had a significant effect on beta-diversity of faecal fungal communities, but total absorbed dose rate category (F_1,111_ = 1.157, R^2^ = 0.010, p = 0.153) and sex (F_1,111_ = 0.823, R^2^ = 0.007, p = 0.911) did not. Burn category effects on faecal community composition were clearer from the fungal community PCA plot (Figure S5) than for the bacterial community PCA (Figure S3); individuals captured in the burnt areas with minimal regrowth tended to appear in the lower left-hand side of the plot (Figure S5). When looking at the samples from yellow-necked mice, differences in clr values for the 12 most abundant fungal genera based on burn category were more pronounced than for bacterial genera (Figure S6). For example, yellow-necked mice trapped in unburnt areas had faecal communities characterised by low *Gelatoporia*, *Pyrenochaetopsis* and *Wickerhamomyces* relative to burnt areas, as well as high *Tritirachium* (Figure S6).

### How do alpha-diversity and beta-diversity of bank vole gut samples vary according to site category and total absorbed dose rate?

Converse to faecal samples, the PERMANOVA analysis showed total absorbed dose rate category had a significant effect on bacterial community beta-diversity of bank vole guts (F_2,138_ = 2.706, R^2^ = 0.036, p = 0.001; Figure S7), as did sex (F_1,138_ = 1.296, R^2^ = 0.009, p = 0.028) and transect line (F_14,138_ = 1.558, R^2^ = 0.146, p = 0.001), whereas site category (i.e. inside or outside the Red Forest) did not (F_1,138_ = 1.221, R^2^ = 0.008, p = 0.091). There were subtle differences evident in the abundance of the 12 most abundant genera across the three total absorbed dose rate categories, of note are the increases in Lachnospiraceae NK4A136, Roseburia, Ruminiclostridium 9, and UBA1819 as radiation dose increased and the decrease in unidentified SVs from the Muribaculaceae family (Figure S8).

The same results were also found for fungal communities of bank vole gut samples, whereby total absorbed dose rate category (F_2,107_ = 2.428, R^2^ = 0.044, p = 0.001; Figure S9) and transect line (F_13,107_ = 1.335, R^2^ = 0.151, p = 0.001) had a significant effect on community beta-diversity, whereas site category did not (F_1,107_ = 1.066, R^2^ = 0.009, p = 0.384). However unlike with bacterial community beta-diversity, sex did not have a significant effect on fungal community beta-diversity (F_2,107_ = 1.174, R^2^ = 0.010, p = 0.142). Again, subtle differences between total absorbed dose rate categories were evident in the 12 most abundant fungal genera, including steady increases in *Arthrobotrys* and *Aspergillus*, and steady decreases in *Candida* and *Wickerhamomyces*, as total absorbed dose rate category increased from low to high (Figure S10).

The partial Mantel test for association between microbial community beta diversity and total absorbed dose rate distance (weighted by geographic distance) indicated a significant relationship for bacterial communities of bank vole guts (r = 0.078, p = 0.047), but the same pattern was not found for fungal communities (r = −0.095, p = 0.866).

There were no significant effects of total absorbed radiation dose rate on alpha-diversity of microbial communities (all p > 0.05; Table S3).

### How do different microbial taxa correlate with the two radiation dose measures?

The association analysis identified seven bacterial families in bank vole gut samples that significantly correlated with total absorbed dose rate (Table S2). All of these had a negative relationship with dose rate with the exception of Lachnospiraceae (Table S2). Two fungal families in bank vole gut samples were significantly correlated with total absorbed dose rate, with Steccherinaceae negatively correlated, and Strophariaceae positively correlated (Table S2). Faecal and gut samples of bank voles showed considerably different fungal and bacterial association patterns (Figures 4 and S11) (note that faecal samples were collected in 2017 and guts, from different animals, in 2018). Fungal and bacterial association patterns between faecal samples from the four small mammal species were also markedly different to one another (Figures 2b and S11b).

**Figure 4.**
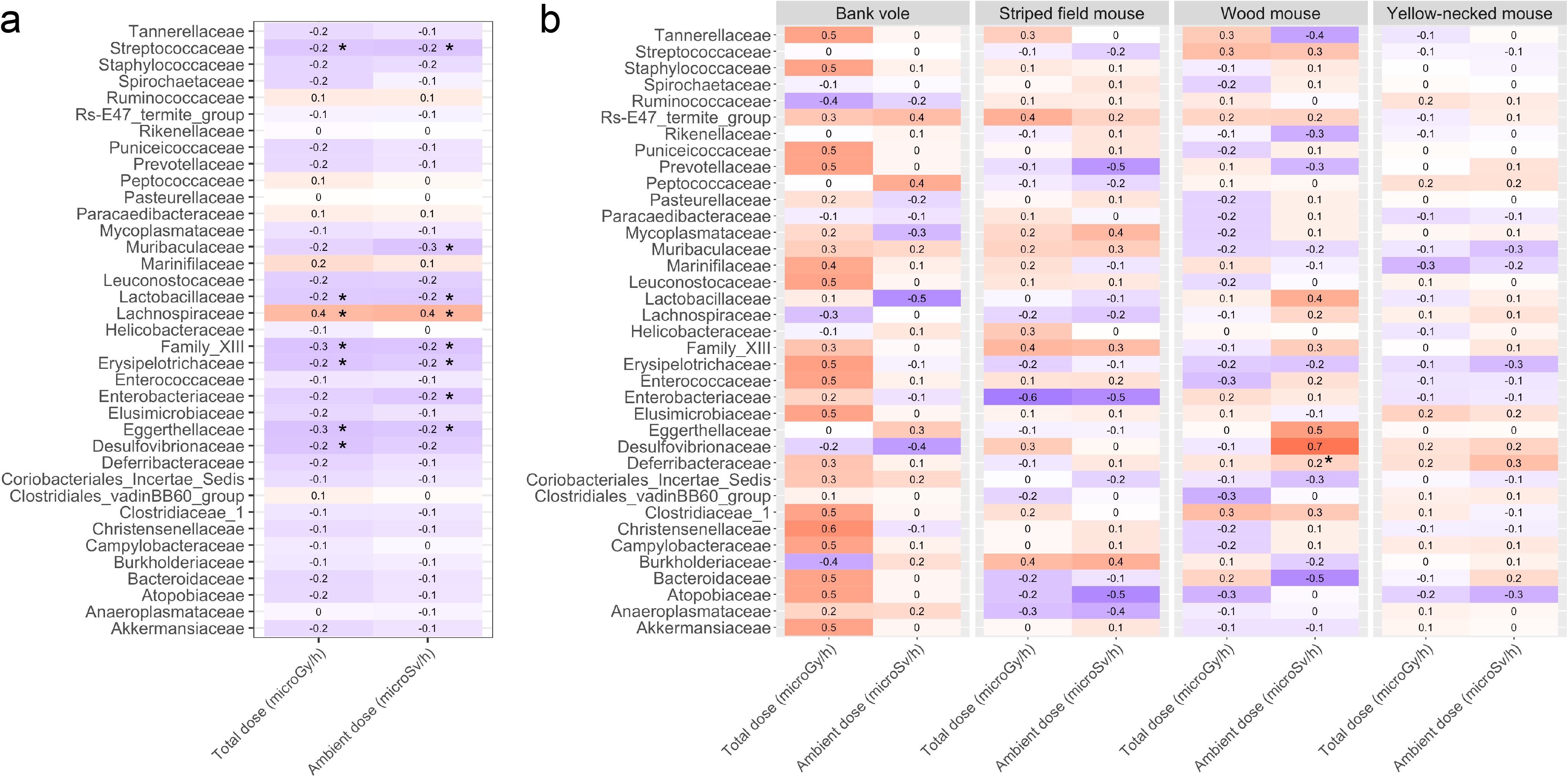
Heatmaps showing correlations between the two radiation dose rate measures (total and ambient) and (a) clr-values of bacterial genera in vole guts and (b) clr-ratios of bacterial genera in faecal samples from four small mammal species. Statistically significant relationships are denoted by *.

### How do Firmicutes: Bacterioidetes ratios vary according to total absorbed dose rate category?

When using clr-transformed data, F:B ratios were less than 0 in vole guts (Figure S12a). When using relative abundance data, voles in the ‘high’ total absorbed dose rate category had slightly higher F:B ratios than those in the ‘low’ and ‘medium’ categories (Figure S12b). The Kruskal-Wallis model indicated a significant effect of total absorbed dose rate category (X^2^ = 7.489, d.f. = 2, p = 0.024), with bank voles in the ‘high’ category exhibiting significantly higher F:B ratios than those in the ‘medium’ category (p = 0.019), but ‘low’ was not significantly different to ‘high’ or ‘medium’ (p > 0.05; Figure S12b). For the faecal sample data, only striped field mice and yellow-necked mice had data for animals in more than one absorbed dose rate category, i.e. medium and high for both. The Kruskal-Wallis analysis was not statistically significant for either striped field mice (X^2^ = 0.012, d.f. = 1, p = 0.911) or yellow-necked mice (X^2^ = 0.019, d.f. = 1, p = 0.896), meaning there were no differences in F:B ratios between the ‘medium’ and ‘high’ categories for these two species (Figure S12c).

## DISCUSSION

In this study we present the first analyses of small mammal faecal and gut microbial communities from the Chernobyl Exclusion Zone for which individual total absorbed dose rates have been estimated. This study also presents the first data from Chernobyl on the fungal component of the gut microbiome, and considers a wider range of species than has previously been studied (previous studies being limited to bank voles ^25,26^). Previous papers have used faecal samples to characterise the gut microbiome ^25,26^, whereas our study provides the first data on the true gut microbiome of Chernobyl bank voles using samples from the distal section of the caecum.

We provide novel evidence that radiation has a small (R^2^ < 0.05), but statistically significant, association with changes in microbial communities of small mammals. We also identified a limited number of taxa with a significant association (Table S2; Figures 4 and S11) with total absorbed dose rate of the host animals. For subsequent discussion of these findings, the relevance of the total absorbed dose rate and its association with specific geographic locations in the CEZ needs to be considered. The total absorbed dose rate is that of the host organism; we have not estimated the radiation exposure of gut microbiota directly. All animals in the ‘high’ total absorbed dose rates (>42 μGy h^−1^) except two bank voles from 2018 were collected from within the Red Forest. Other studies of radiation effects in CEZ wildlife, including the microbiome studies of Lavrinienko et al. ^25,26^, also have their most contaminated sampling sites within the Red Forest. The Red Forest is an area of naturally poor habitat quality where soil and water conditions do not favour high biological diversity. The forest was also severely damaged in 1986 as a consequence of the accident at the Chernobyl Nuclear Power Plant and has not fully recovered. Furthermore, some of our 2017 Red Forest sampling sites were showing signs of fire damage from a large fire in July 2016. Any study that uses the Red Forest as a location for radiation effects studies on wildlife needs to consider the historical impacts of radiation and other stressors (e.g. wildfires) on this area of the CEZ ^63^.

Geographical location is known to affect bacterial community composition ^41,64^, and here we find the same, whereby grid/transect line are significant predictors of bacterial beta-diversity. We also provide novel evidence that geography affects fungal community composition. The Mantel test shows that bacterial community composition of bank vole guts varied predictably with total absorbed dose rate when taking geographical variation into account (although the same was not true for fungal communities). Furthermore, the PERMANOVA analysis of bank vole gut microbiome indicated that total absorbed dose rate category was a significant predictor of both bacterial and fungal beta-diversity, whereas site category (inside or outside the Red Forest) was not. Together, these results indicate that differences in microbiome composition exhibited in bank vole guts were a result of radiation exposure, rather than the confounding effects of geography or habitat type. However, given that microbes are actually highly resilient to radiation exposure ^65,66^, environmental radiation exposure at sites such as Chernobyl is unlikely to affect the microbiome directly. It is more likely that co-correlating factors are driving observed relationships between radiation and gut microbiome ^67^. For example, radiation exposure that causes changes in habitat quality, food availability or a host physiological response may all influence the gut microbiome ^68–70^.

The gut communities of bank voles showed similar changes in composition in response to both ambient and total dose radiation measures, although relationships were generally less strong for ambient dose compared with total dose (Figures 4 and S11). This may be due to differences in the way that individual dose rates are assigned; every individual from a site is assigned the same dose rate whereas total absorbed dose rate is calculated on an individual basis. The bacterial genera identified here (Figure 4; Table S2) may serve as useful bioindicators for radiation exposure in mammals, although more work is required to determine if these patterns are consistent across different host species. Our data from faecal samples indicate that the relationship between the small mammal microbiomes and total absorbed dose rate of the host may vary from species to species (Figures 4 and S11), although there were relatively few significant relationships (identified by * on Figure 4) between radiation dose and clr values for individual genera. However, these heatmaps indicate that faecal sample communities exhibited considerably different results for the two dose measures, suggesting that faecal samples and/or ambient dose measures are not a reliable method of characterising microbiome changes in response to radiation exposure.

The relationships between microbial families and radiation exposure were considerably different between gut and faecal samples for bank voles. This may be because different taxa are being excreted to those that are retained in the gut. It has previously been shown that a number of host species have significantly different communities associated with the gut and faeces ^71,72^. As such, faecal samples may not directly reflect responses of gut communities to radiation exposure or any other stressor. However, it is worth noting that bank vole gut samples were collected in 2018 from across the CEZ, whereas the faeces samples collected in 2017 were all from inside the Red Forest (including from a number of sites that had been recently burnt), which may also be influencing the observed differences between the gut and faecal samples.

Firmicutes and Bacteroidetes are the most abundent phyla within the microbiome of small mammals; Firmicutes have been linked to processes such as the generation of metabolites, fat storage, angiogenesis and immune system maturation ^10^. Lavrinienko et al. ^25^ found a two-fold increase in F:B ratios in bank vole faeces from areas of elevated radionuclide contamination in the CEZ (these sites would span our ‘medium’ and ‘high’ total absorbed dose rate categories) compared with areas of lower contamination in the CEZ and sites close to Kiev. The authors attribute the two-fold increase in F:B ratios to potential changes in diet arising from reduced arthropod densities in their higher contamination areas of the Chernobyl Exclusion Zone (referring to earlier work of Møller & Mousseau ^73^, the findings of which have been contested ^30,74,75^) and/or an active increase in the consumption of plant based foods. Indeed, F:B ratios in faeces have previously been used as a marker of changes in diet ^76^. However, the authors also state that the bank vole diet is normally dominated by plant material, with only occasional consumption of invertebrates. Consequently, the effect of any reduction in arthropod consumption on the bank vole faecal microbiome F:B ratios would likely be minimal. In the present study, we found no evidence of altered F:B ratios in bank vole gut samples based on total absorbed radiation dose rate category suggesting similar bank vole diets across our study locations, including inside and outside the Red Forest.

Our results suggest that bacterial communities are more influenced by total absorbed dose rate than fungal communities. For bank vole guts, the partial Mantel tests were not significant for fungi (p > 0.80) but were for bacteria (p < 0.05). Fewer fungal families than bacterial families were significantly associated with total absorbed dose rate (Table S2). In addition, fungi and bacteria appeared to display opposing responses to radiation. For example, the heatmaps suggest that family-level associations with total absorbed dose rate in bank vole guts were mostly positive for fungi (Figure S11), but mostly negative for bacteria (Figure 4). Together, our results suggest that changes in host-associated fungal communities may be less associated with radiation exposure than changes in bacterial communities (Table S2).

To our knowledge, we present here the first demonstration that host species is a significant predictor of fungal community composition in ground-dwelling small mammal populations; fungal community compositions are an under-explored aspect of host-associated microbiomes in general ^9,77^. In agreement with previous studies on a range of species ^78–80^, including small mammals ^4^, we also find host species to be a significant predictor of bacterial community composition. We found no effect of sex on bacterial or fungal communities of faecal samples from any host species, or on fungal communities of bank vole guts. However, we did find that sex had a significant effect on bacterial community composition of bank vole guts. Previous studies have found mixed effects of sex on microbiome composition of small mammals ^4,10,25^.

## Conclusions

Using a range of statistical analyses, we identify a number of significant associations between total absorbed radiation dose and changes in microbiome, particularly for the bacterial component. For bank vole gut data, these results were robust against confounding factors including geographic variation and habitat type. However, the overall evidence for a significant impact of radiation on the fungal component of the microbiome was limited. Furthermore, contrary to the findings of the only previous published study of small mammal microbiome in the CEZ ^25,26^, we did not see any significant effect on the F:B ratio with absorbed dose rate. We also provide evidence that faecal samples are not reliable for examining microbiome changes in response to radiation, and that total absorbed radiation doses provide considerably more accurate results than ambient dose measures.

We suggest that, given the importance of the microbiome to host health and the limited studies on microbiome (especially fungal microbiome and gut microbiome) in wild animals, further studies of the microbiome response to radiation and other factors within the CEZ should be undertaken. In particular, there are outstanding questions around whether host microbiomes are directly affected by radiation exposure, or rather mediated by some mechanism of host physiology. For this, it is important to establish directionality; i.e. whether the host microbiome alters host physiology in response to radiation exposure, or vice versa. Furthermore, changes in diet, resulting from impacts of radiation on the ecosystem (e.g. in an area such as the Red Forest) rather than the host, may also be expected to affect the gut microbiome. More work is required to understand the mechanisms that are driving changes in host microbiomes of wildlife in general, and the implications of this for host function and fitness.

## Supporting information

Radioecology supplementary data

Figure S11 high definition

## ACKNOWLEDGEMENTS

The work described in this paper was conducted within the TREE (https://tree.ceh.ac.uk/) and RED FIRE (https://www.ceh.ac.uk/redfire) projects. TREE was funded by the Natural Environment Research Council (NERC), Radioactive Waste Management Ltd. and the Environment Agency as part of the RATE Programme; RED FIRE was a NERC Urgency Grant. The study was undertaken in line with ethical approval obtained from the University of Salford. For the 2017 study, the GMS 310 core gamma logger was kindly loaned by John Caunt Scientific Ltd.

## STATEMENT OF AUTHORSHIP

MDW, NAB & SG designed the study and undertook sample collection (along with RF, JAJ, CLB, EP & LW); SG characterised sites; NAB and RF live-monitored small mammals; REA conducted the DNA extraction, molecular work and statistical analysis; REA, NAB and MDW wrote the paper; all authors revised and approved the final manuscript.

## DATA ACCESSIBILITY STATEMENT

Sequence data are available from the NCBI SRA database under project numbers PRJNA594002 and PRJNA592322.

