## Supplementary material for "Impacts of radiation on the bacterial and fungal microbiome of small mammals in the Chernobyl Exclusion Zone"

**Table S1**

Sample sizes of host species used in the study, and associated sex, total absorbed dose rate, burn and site categories for these samples. Animals estimated to receive total absorbed dose rates of <4 μGy $h^{-1}$ were assigned 'low', those with estimated dose rates of 4-42 μGy $h^{-1}$ assigned 'medium', and those >42 μGy $h^{-1}$ assigned to the 'high' category. All samples from 2017 were collected within the Red Forest but from areas that had experienced different degrees of damage from forest fires. Samples from 2018 were either collected within or outside the Red Forest.

| Sampling information | | | | Sex | | | Total absorbed dose category | | | Burn category | | | Site category | |
|---|---|---|---|---|---|---|---|---|---|---|---|---|---|---|
| Year | Sample type | Host species | Total n | Female | Male | Unidentified | Low | Medium | High | Burnt (regrowth) | Burnt (minimal regrowth) | Unburnt | Inside Red Forest | Outside Red Forest |
| 2017 | Faeces | Bank vole | 22 | 5 | 16 | 1 | 0 | 0 | 22 | 3 | 1 | 18 | All | - |
| 2017 | Faeces | Striped field mouse | 29 | 10 | 18 | 1 | 0 | 7 | 22 | 15 | 0 | 14 | All | - |
| 2017 | Faeces | Wood mouse | 27 | 6 | 20 | 1 | 0 | 0 | 27 | 2 | 25 | 0 | All | - |
| 2017 | Faeces | Yellow-necked mouse | 58 | 28 | 29 | 1 | 0 | 7 | 51 | 19 | 17 | 22 | All | - |
| 2018 | Gut (caecum) | Bank vole | 142 | 59 | 83 | 0 | 42 | 64 | 36 | - | - | - | 34 | 108 |

### RESULTS

#### *Differences in community composition between host species and sample types*

Wood mice and yellow-necked mice had a similar bacterial community composition to one another (Figure 1a), with the exception of the family Helicobacteraceae ($Z = 2.982$, $p = 0.023$; Figure 1c). Striped field mice had a different bacterial community structure to the other two mice species (Figure 1a), characterised by higher Enterobacteriaceae and lower Eggerthellaceae compared with both of the other mice species, as well as lower Enterococcaceae than wood mice; lower Burkholderiaceae than yellow-necked mice; and higher Helicobacteraceae, Bacteroidaceae and Paracaedibacteraceae than yellow-necked mice (all $p < 0.001$; Figure 1c). Bank voles had considerably different faecal bacterial community composition to the four mice species; bank voles were comparatively dominated by Paracaedibacteraceae, but lower in Bacteroidaceae, Enterococcaceae, Deferribacteraceae and Burkholderiaceae compared with all three mice species, as well as lower Eggerthellaceae compared to yellow-necked and wood mice, and lower Enterobacteriaceae than striped mice (all $p < 0.001$; Figure 1c). Bank vole guts had significantly lower Muribaculaceae, Paracaedibacteraceae and Coriobacteriales incertae sedis compared with bank vole faecal samples, and significantly higher Clostridiales vadinBB60 (all $p < 0.001$; Figure 1c). Only five of the12 most abundant families identified from clr-transformed data were also identified as the 12 most abundant using relative abundance data (Figures 1c and S1). For those families in common (Bacteroidaceae, Enterobacteriaceae, Helicobacteraceae, Muribaculaceae, Ruminococcaceae), patterns were broadly similar between the two data normalisation techniques (Figures 1c and S1).

Fungal communities of faecal samples from the three mice species overlapped considerably (Figure 1b), with the only statistically significant difference in the top 12 most abundant families being lower Tritirachiaceae in wood mice relative to the other two mice species (all $p < 0.001$; Figure 1d). Bank vole faecal samples had higher Psathyrellaceae than all three mice species; higher Tritirachiaceae than wood mice; and lower Aspergillaceae than wood mice and yellow-necked mice (all $p < 0.001$; Figure 1d). Differences in fungal community composition between the two sample types from bank voles were higher Phaffomycetaceae and Psathyrellaceae in gut samples, and higher Pleosporaceae in faecal samples (all $p < 0.001$; Figure 1d). Ten families were represented in both the clr-transformed and relative abundance data (Aspergillaceae, Debaryomycetaceae, Hypocreaceae, Meruliaceae, Phaffomycetaceae, Pleosporaceae, Psathyrellaceae, Saccharomycetales fam incertae sedis, Saccharomycodaceae, Trichocomaceae), again with broadly similar patterns between the two normalisation methods (Figure 1d and S2).

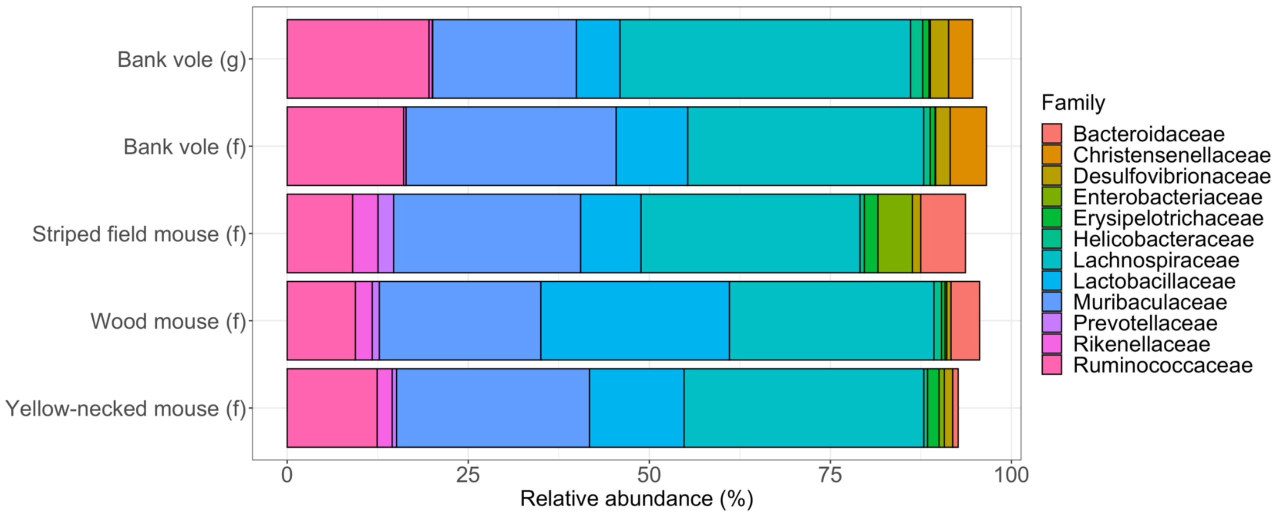


**Figure S1**

Relative abundance of the 12 most abundant bacterial families across the five main sampling groups in the study.

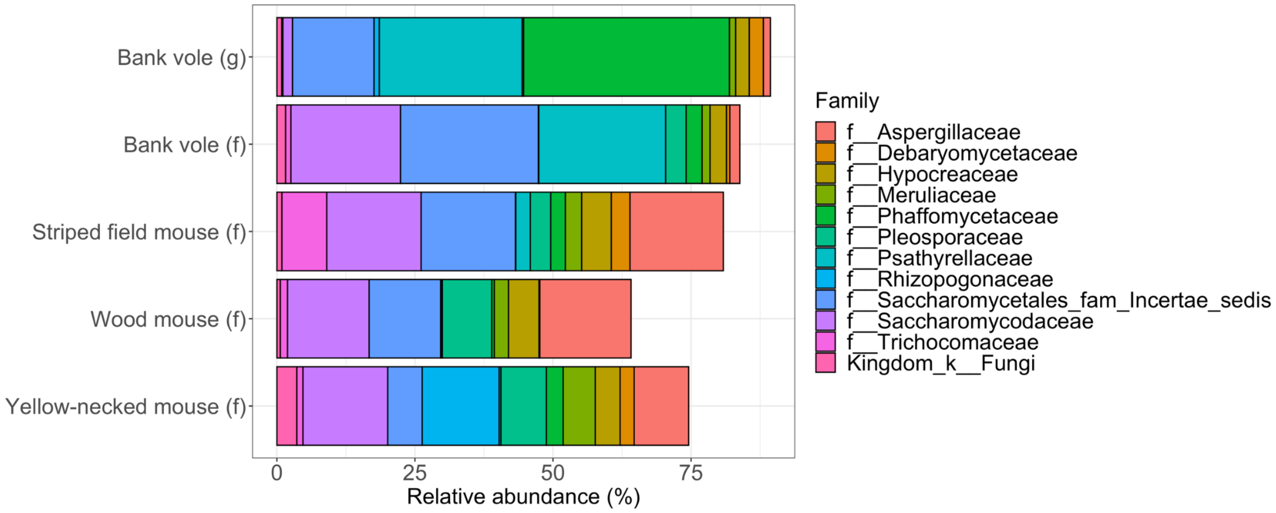


**Figure S2**

Relative abundance of the 12 most abundant fungal families across the five main sampling groups in the study. Note “Kingdom_k_Fungi” refers only to fungi unidentified at the kingdom level.

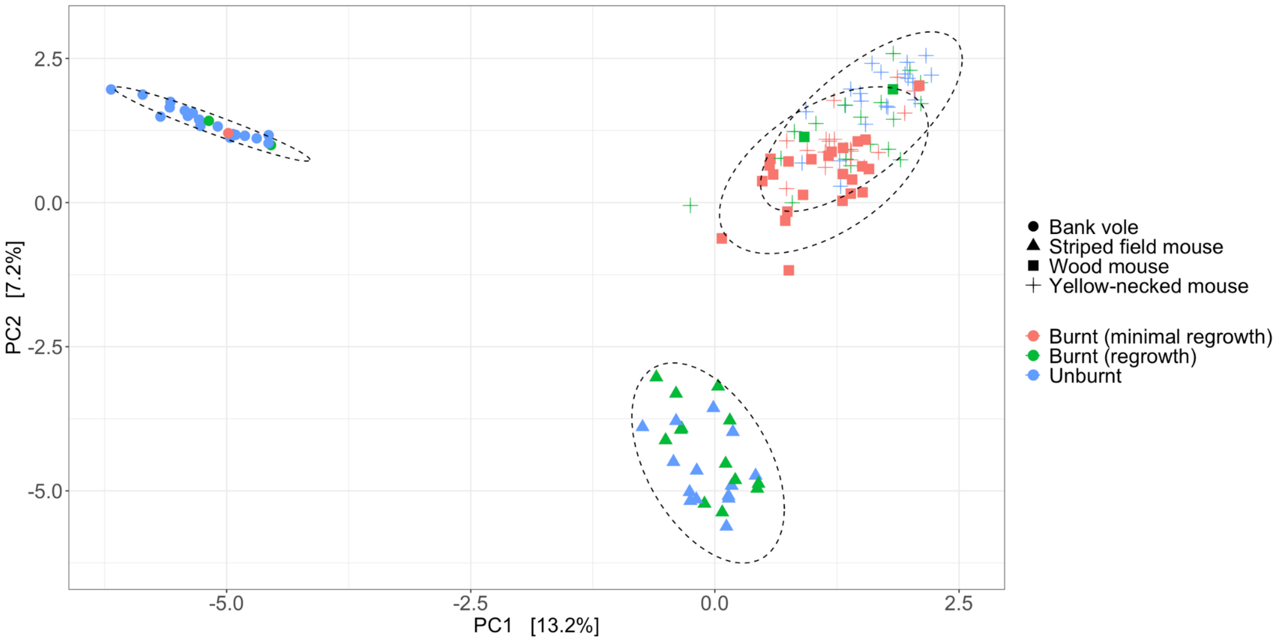


**Figure S3**

PCA plot showing Euclidean distances of clr-transformed bacterial communities associated with faecal samples from four small mammal species in the Chernobyl Exclusion Zone, coloured by burn category in which the animals were captured.

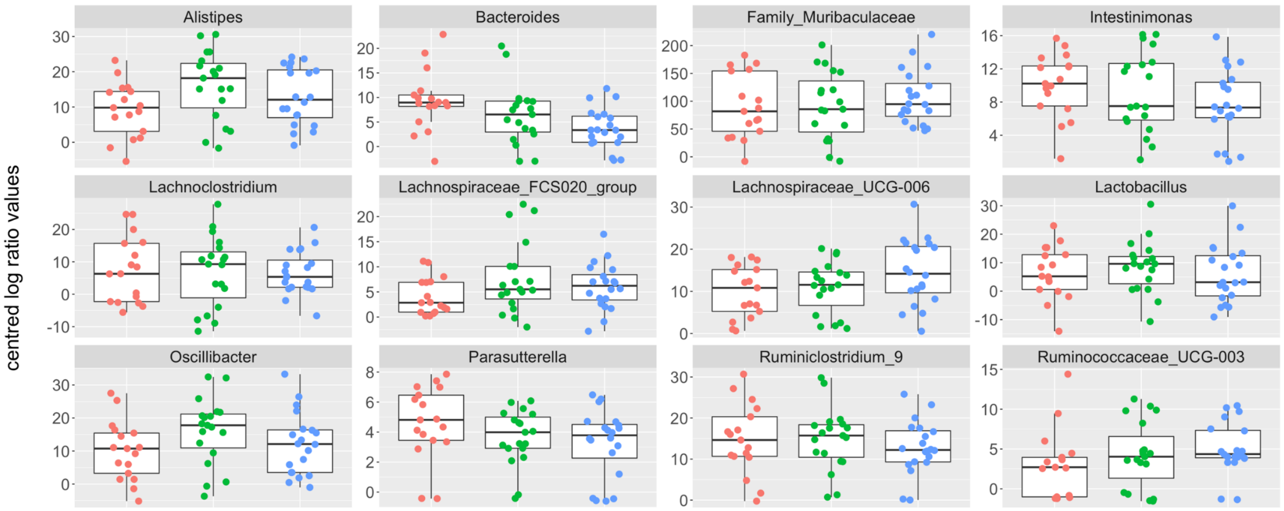


**Figure S4**

Jitter plots of clr values of the 12 most abundant bacterial genera in faecal samples of yellow-necked mice in three burn categories in the Chernobyl Exclusion Zone; burnt with minimal regrowth shown in red, burnt with regrowth shown in green, and unburnt shown in blue.

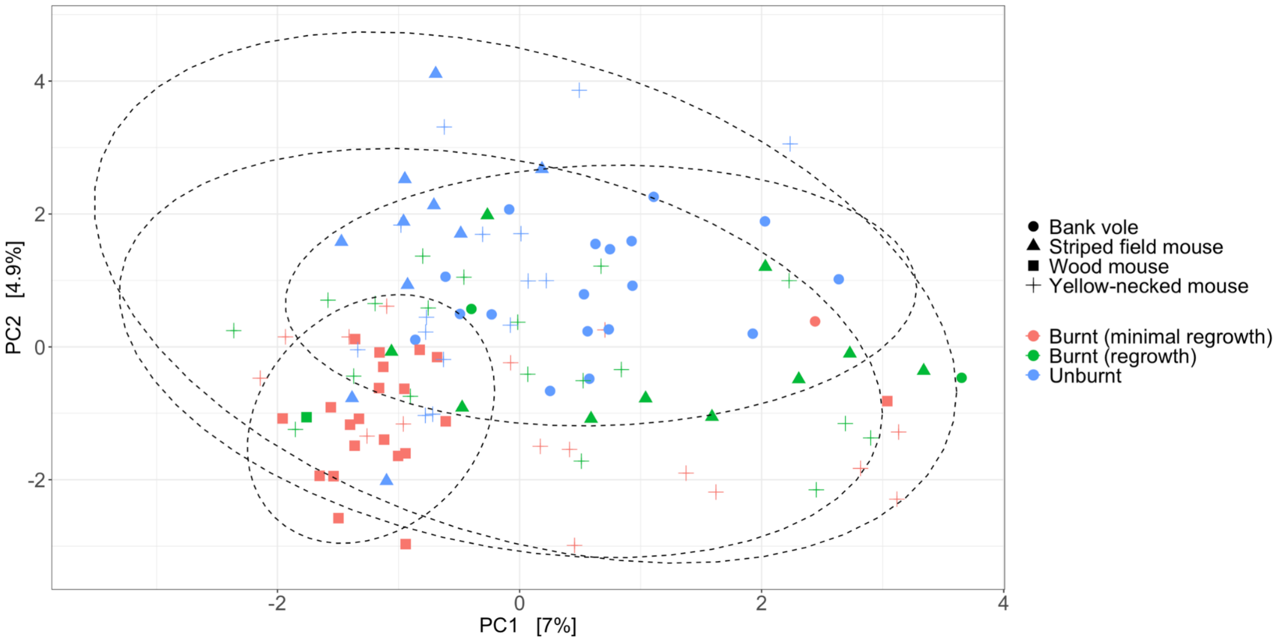


**Figure S5**

PCA plot showing Euclidean distances of clr-transformed fungal communities associated with faecal samples from four small mammal species in the Chernobyl Exclusion Zone, coloured by burn category in which the animals were captured.

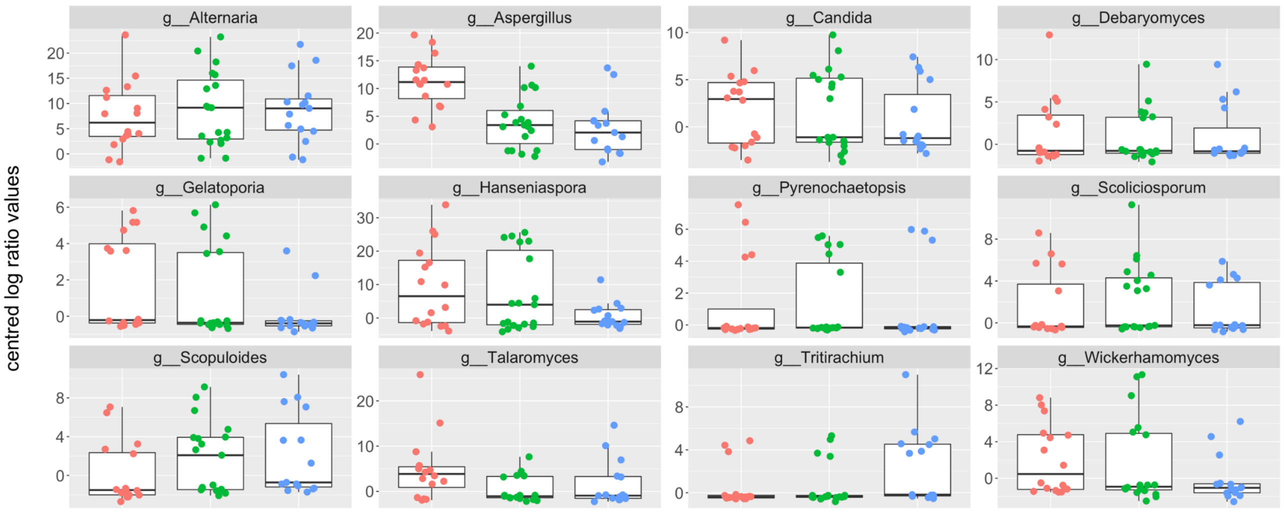


**Figure S6**

Jitter plots of clr values of the 12 most abundant fungal genera in faecal samples of yellow-necked mice in three burn categories in the Chernobyl Exclusion Zone; burnt with minimal regrowth shown in red, burnt with regrowth shown in green, and unburnt shown in blue.

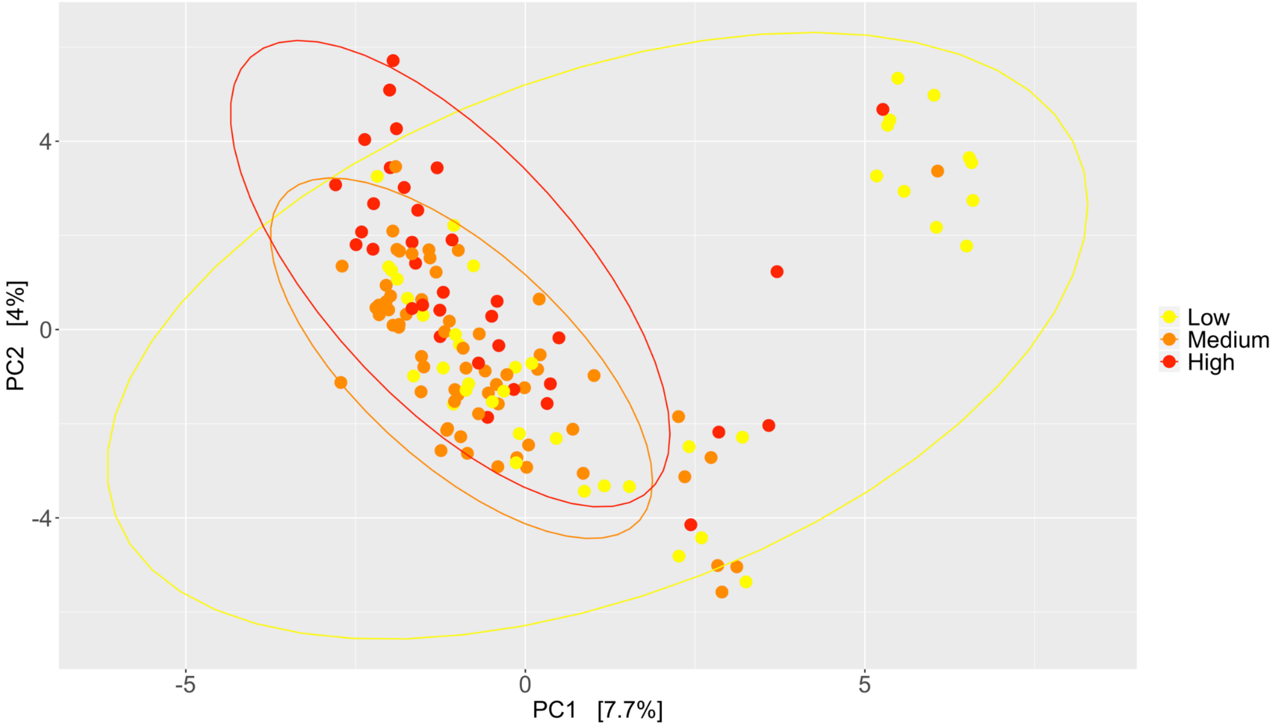


**Figure S7**

PCA plot of bank vole gut bacterial communities according to total absorbed dose rate category (low <4 microGy/h, medium = 4-42 microGy/h, and high > 42 microGy/h.

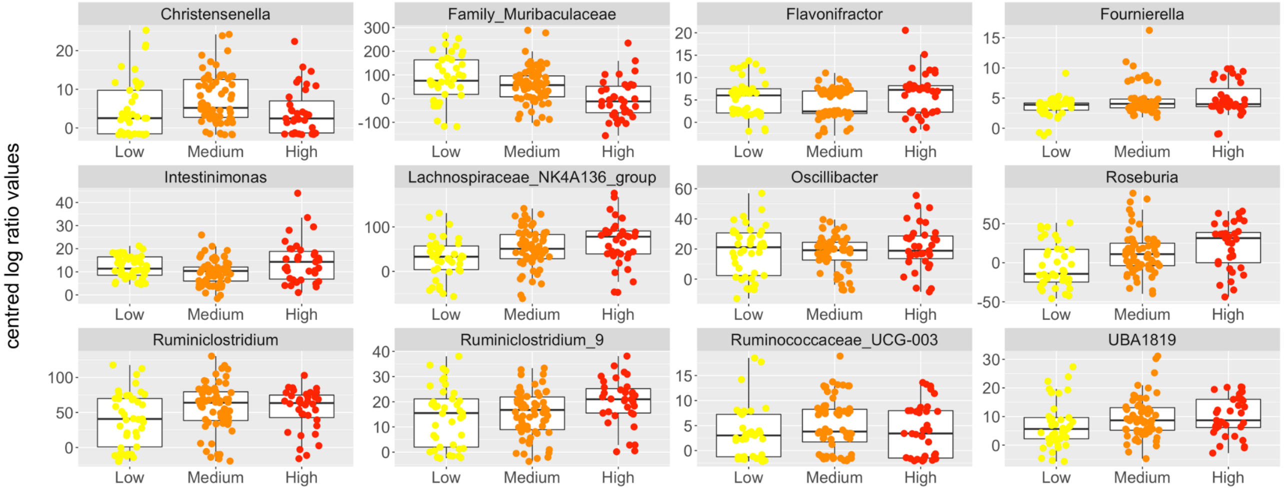


**Figure S8**

Jitter plots of clr values of the 12 most abundant bacterial genera in bank vole gut bacterial communities according to total absorbed dose rate category (low <4 microGy/h, medium = 4-42 microGy/h, and high > 42 microGy/h).

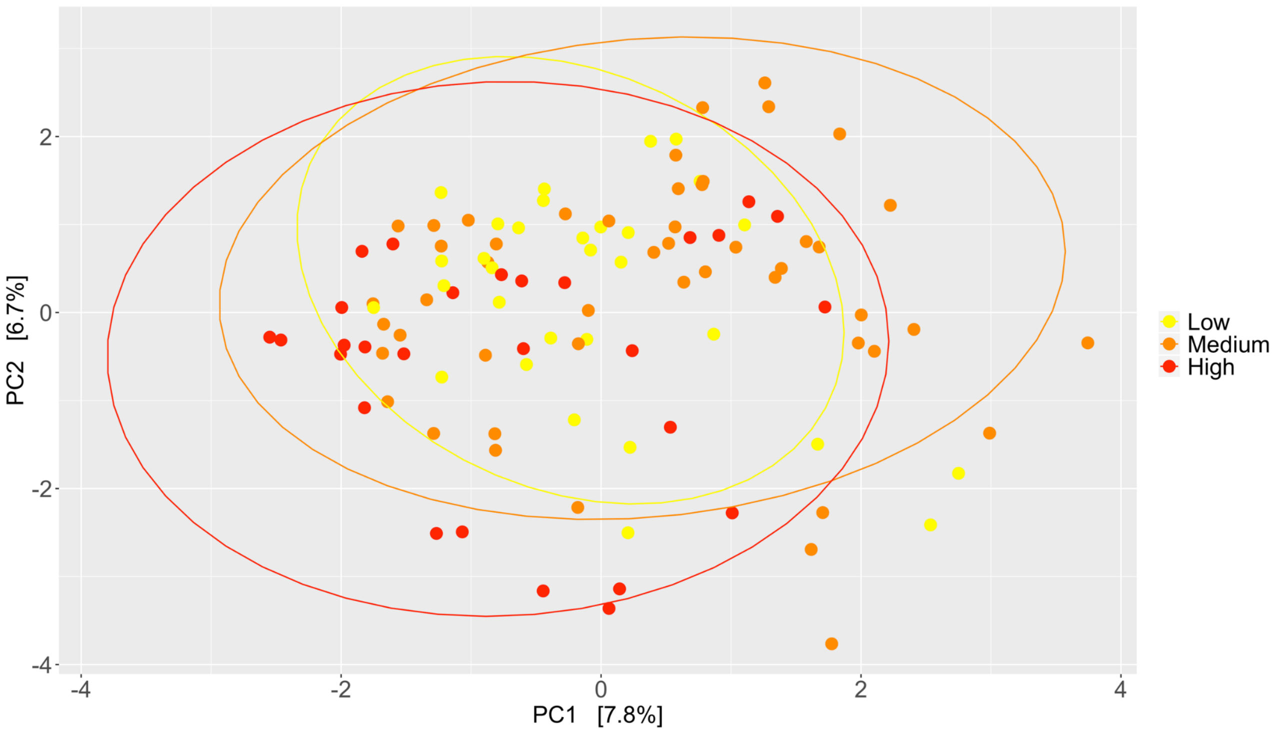


**Figure S9**

PCA plot of bank vole gut fungal communities according to total absorbed dose rate category (low <4 microGy/h, medium = 4-42 microGy/h, and high > 42 microGy/h)

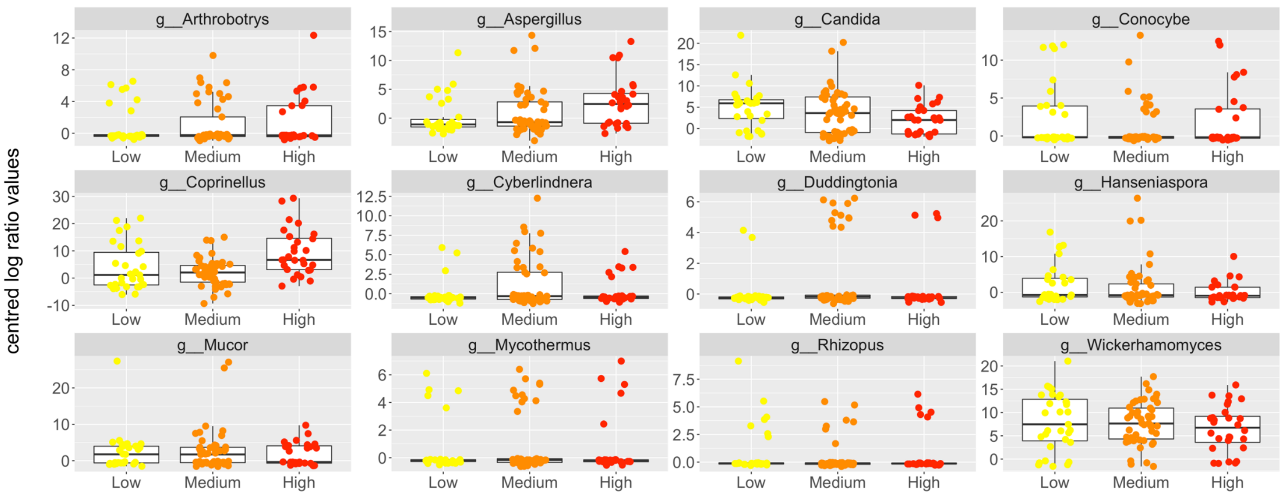


**Figure S10**

Jitter plots of clr values of the 12 most abundant fungal genera in bank vole gut bacterial communities according to total absorbed dose rate category (low <4 microGy/h, medium = 4-42 microGy/h, and high > 42 microGy/h).

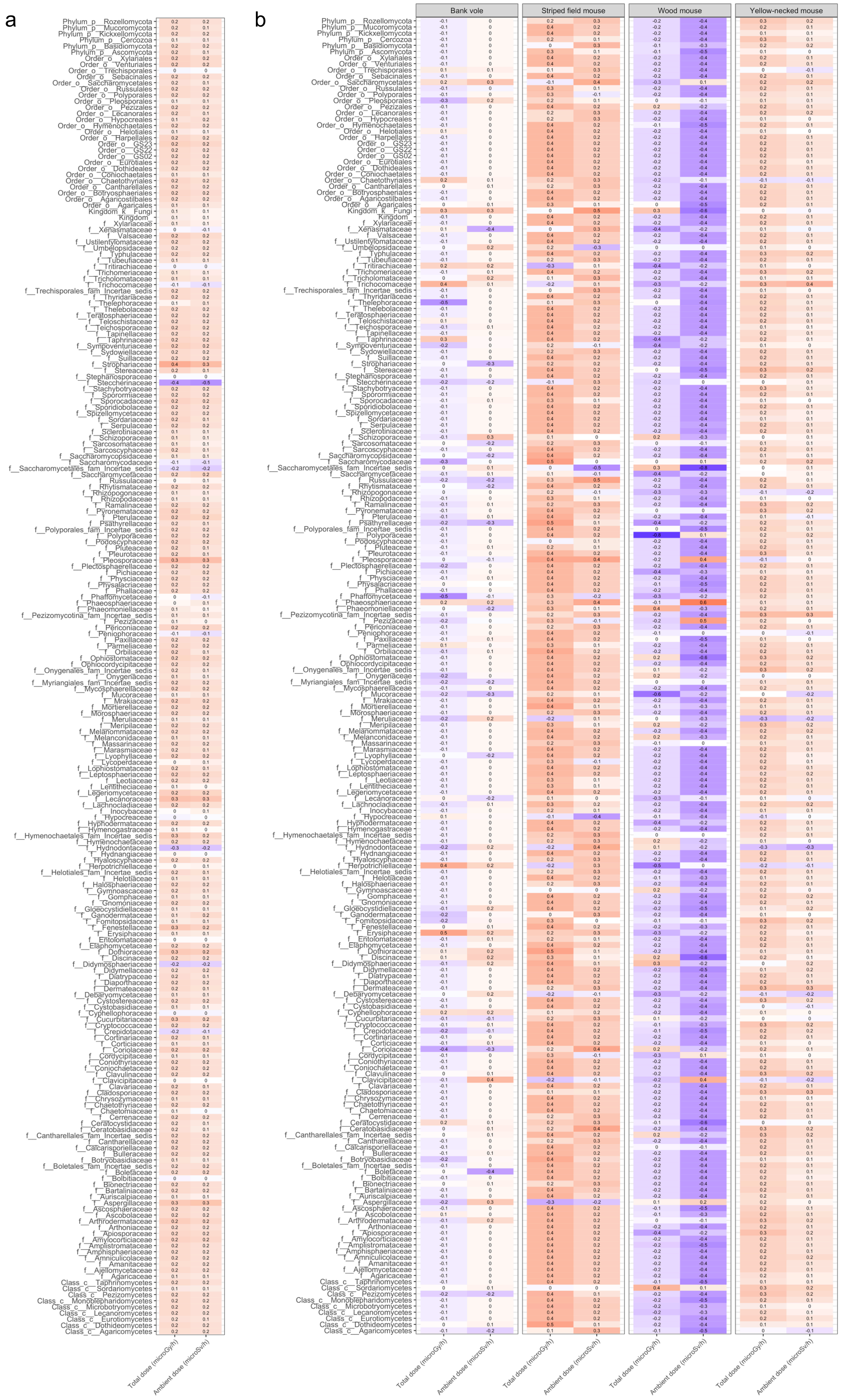

**Figure S11**

Correlations between the two radiation dose measures (total absorbed and ambient dose rates) and (a) clr-ratios of fungal genera in vole guts and (b) clr-ratios of fungal genera in faecal samples from four small mammal species (see additional figure provided as a supplementary file for easier viewing).

**Table S2**

Microbial families from the guts of bank voles significantly correlated with total absorbed dose rate, as identified through association analysis.

| Kingdom | Family | Correlation (r) | Adjusted p value |
|---|---|---|---|
| Bacteria | Desulfovibrionaceae | -0.247 | 0.030 |
| Bacteria | Eggerthellaceae | -0.259 | 0.030 |
| Bacteria | Erysipelotrichaceae | -0.245 | 0.030 |
| Bacteria | Family_XIII | -0.258 | 0.030 |
| Bacteria | Lachnospiraceae | 0.360 | <0.001 |
| Bacteria | Lactobacillaceae | -0.221 | 0.047 |
| Bacteria | Streptococcaceae | -0.237 | 0.033 |
| Fungi | Steccherinaceae | -0.428 | 0.009 |
| Fungi | Strophariaceae | 0.418 | 0.010 |

**Table S3**

Results of Spearman's rank correlations between microbial alpha-diversity of bank vole guts and total absorbed dose rate.

| Diversity measure | Microbial kingdom | r | p value |
|---|---|---|---|
| Richness | Bacteria | -0.054 | 0.528 |
| Evenness | Bacteria | 0.006 | 0.942 |
| Richness | Fungi | -0.143 | 0.185 |
| Evenness | Fungi | -0.134 | 0.217 |

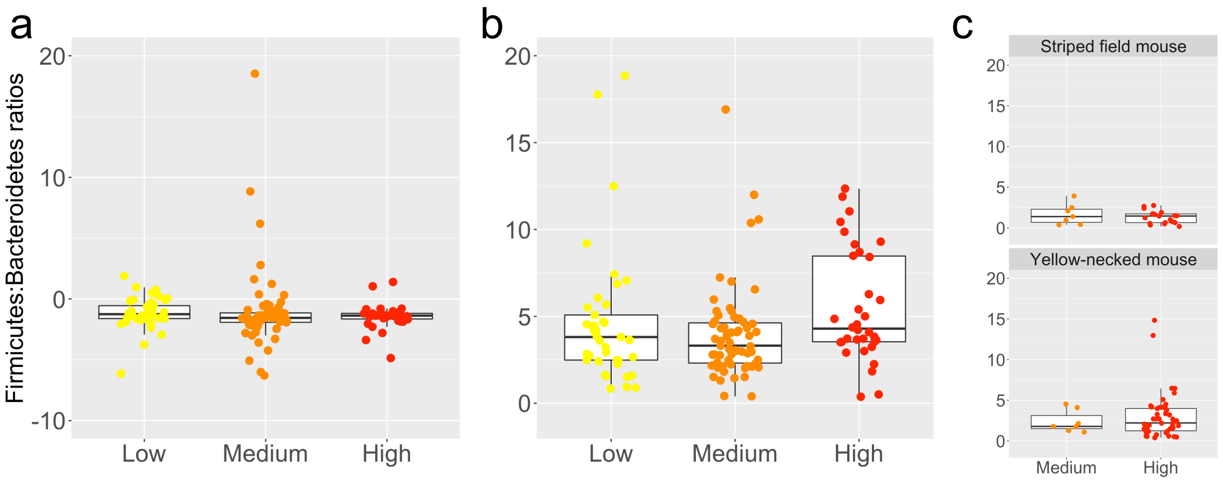


**Figure S12**

Firmicutes: Bacteroidetes ratios according to total absorbed dose rate category based on a) clr values of bank vole guts; b) relative abundance values of gut samples from bank voles; and c) relative abundance values of faecal samples from four small mammal species.
