## Supplementary figures and images for "Impacts of radiation on the bacterial and fungal microbiome of small mammals in the Chernobyl Exclusion Zone"

### Figure S11 high definition

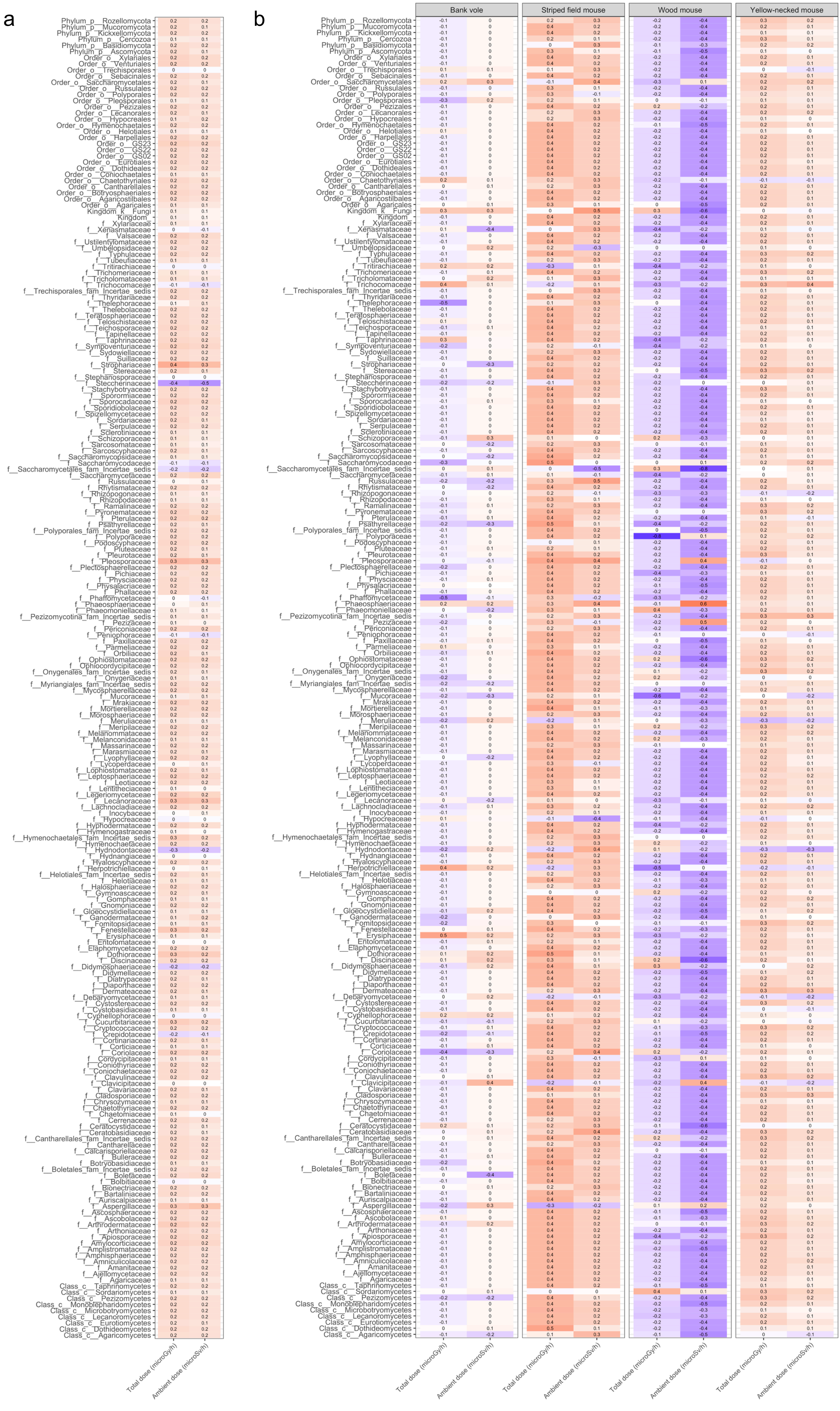
